# Magnesium Sulfate Attenuates Lethality and Oxidative Damage Induced by Different Models of Hypoxia in Mice

**DOI:** 10.1101/2020.05.22.111435

**Authors:** Hamidreza Mohammadi, Amir Shamshirian, Shafagh Eslami, Danial Shamshirian, Mohammad Ali Ebrahimzadeh

## Abstract

Mg^2+^ is an important cation in our body. It is an essential co-factor for many enzymes. Despite many works, nothing is known about the protective effects of MgSO_4_ against hypoxia-induced lethality and oxidative damage in brain mitochondria. In this study, antihypoxic and antioxidative activities of MgSO_4_ were evaluated by three experimental models of induced-hypoxia (asphyctic, haemic, and circulatory) in mice. Mitochondria protective effects of MgSO_4_ were evaluated in mice brain after induction of different models of hypoxia. Antihypoxic activity was especially pronounced in asphyctic hypoxia where MgSO_4_ at dose 600 mg/kg showed the same activity as phenytoin which used as a positive control (*P<* 0.001). In the haemic model, MgSO_4_ at all used doses significantly prolonged latency of death. In circulatory hypoxia, MgSO_4_ (600 mg/kg) doubles the survival time. MgSO_4_ significantly decreased Lipid peroxidation, protein carbonyl, and improved mitochondrial function and glutathione content in brain mitochondria compared to control groups. The results obtained in this study showed that MgSO_4_ administration has protective effects against lethality induced by different models of hypoxia and improves brain mitochondria oxidative damage.

## 1. Introduction

The cation Mg^2+^ has an important role in the intracellular process regulation. It is necessary for ATP activity containing calcium current across and within cell membranes for tissues. Moreover, Mg^2+^ is existing in more than 300 enzymatic systems through antioxidative properties and is the fourth most cation in our body (1, 2). Mg^2+^ is an essential cofactor for the activity of kinases and effects on oxidative phosphorylation, DNA synthesis, lipid hydrolysis, and rates of glycolysis (3). Subsequently, Mg^2+^ is compulsory in all relating ATP reactions, and variations in this ion may regulate oxidative phosphorylation and substrate in the myocardial (4). Cardiovascular disease has been associated with diminished Mg^2+^ levels and a pivotal relationship may be present between mortality from non-occlusive ischemic heart disease and decreased cardiac Mg^2+^ (4, 5). Myocardial damage consequence of oxidative stress in the heart cells can occur in several conduction problems such as hypoxia. There is evidence that hypoxia reduces cellular ATP, depletes mitochondrial energy, elevates ADP/ATP ratio mainly in the heart muscle cells, and induces oxidative stress (6). The increased generation of ROS can cause a decrease in the rate of ATP synthesis in mitochondria because of the loss of cytochrome oxidase (Cox) activity. Cox is a chain in the mitochondrial respiratory process, which generates ATP by oxidative phosphorylation. The inhibition of Cox appears to be the main reason for the ATP depletion. A reduced capacity of Cox will cause an increase in ROS production and decrease ATP synthesis (4, 5, 7).

The imbalance between the supply of oxygen and its demand determines organ hypoxia. It occurs especially in ischemia and heart diseases and leads to numerous deleterious effects and finally resulting in death (8). Hypoxia causes oxidative stress involving the production of reactive oxygen species (5, 9). It has proven that the compounds with antioxidant activity can scavenge ROS and able to exhibit anti-hypoxic property (10).

MgSO_4_ is a well-known cation that is vital for cell viability. To the best of our knowledge, there is no report about the protective effects of this ion against lethality and mitochondria oxidative brain damage induced by hypoxia. The present work aimed to determine the anti-hypoxic activities and neuroprotective effects of MgSO_4_ to understand a possible mechanism of its action in hypoxia-induced lethality and brain mitochondria oxidative damage protection induced by different models of hypoxia.

## 2. Material and methods

### 2.1. Chemicals

All chemical agents were prepared from Sigma-Aldrich Company and were used in this study.

### 2.2. Animals

Male Swiss albino mice (21 ± 3g) were housed in polypropylene cages at 25 ± 1°C and 45-55% relative humidity, with a 12h light: 12h dark cycle (lights on at 7 a.m.). The animals had free access to standard pellet and water and libitum. Experiments were conducted between 8:00 and 15:00 h. All the experimental procedures were conducted following the NIH guidelines of the Laboratory Animal Care and Use. The Institutional Animal Ethical Committee of Mazandaran University of Medical Sciences also approved the experimental protocol (with ethics number: IR.MAZUMS.REC.93.1001).

### 2.3. Animals treatment

208 male mice were randomly divided into 26 groups (eights in each). 13 groups were used for hypoxia-induced methods and 13 groups were used for evaluation of brain mitochondria oxidative stress biomarkers after induction of different hypoxia methods.

### 2.4. Asphyctic Hypoxia

The animals were subjected to hypoxia by putting them individually in a tightly closed 300 ml glass container which was placed underwater in an aquarium of 25°C. The animals had convulsions and died from hypoxia. The latencies for death were recorded. The animals died approximately 1.5-2 min following convulsions. Mice received single i.p. injections of 300, 400, and 500 mg/kg doses of MgSO_4_ or phenytoin (50 mg/kg) as 30 min before they were subjected to hypoxia. The Control group was treated with normal saline (11).

### 2.5. Haemic Hypoxia

Forty mice were divided into five groups each containing eight mice. Mice were injected with sodium nitrite (NaNO_2_) in dose 360 mg/kg i.p. thirty minutes after the i.p. administration of 400, 500, or 600 mg/kg doses of MgSO_4_. The survival time (in min) for each animal is defined as time, measured from the induction of the hypoxia, caused by the introduction of the hypoxia, until death. The control group was treated with normal saline.

### 2.6. Circulatory Hypoxia

Forty mice were divided into five groups each containing eight mice. The groups were treated with normal saline. Thirty minutes after *i.p.* administration of 400, 500, and 600 mg/kg doses of MgSO_4_, NaF (150 mg/kg) was applied *i.p.* to mice and anti-hypoxic activity was estimated in minutes as the latent time of evidence of hypoxia (12).

### 2.7. Preparation of the brain mitochondria

All animals were sacrificed by cervical decapitation after seven minutes (in haemic and circulatory models) and 25 minutes in the asphyctic model based on our above results. Then, brain tissues were homogenized and mitochondria were extracted from mice brain using differential centrifugation. Tris buffer (0.05 M Tris-HCl, 20 mm KCl, 0.25 M sucrose, 2.0 mM MgCl_2_, and 1.0 mM Na_2_HPO_4_, pH= 7.4) was used for suspending of the final mitochondrial pellets. For ROS production assessment, mitochondria were suspended in respiration buffer (0.32 mM sucrose, 20 mM Mops, 10 mM Tris, 0.5 mM MgCl_2_, 50 μM EGTA, 0.1 mM KH_2_PO_4_ and 5 mM sodium succinate). All procedures were done at 4 °C (13).

### 2.8. Protein Concentration

The Coomassie blue protein binding method (14) was used for the measurement of mitochondria protein content in the samples.

### 2.9. Lipid peroxidation (LPO) measurement

MDA malondialdehyde content was measured by the Zhang *et al.* method (15). Briefly, phosphoric acid (0.25 mL, 0.05 M) was added to mitochondrial fractions (0.2 mL, 0.5 mg protein/ml) and then thiobarbituric acid (TBA) (0.3 mL 0.2% was added to them. Afterward, the microtubes were located (30 min) in a boiling water bath. In the end, the samples were placed to an ice-bath and n-butanol (0.4 ml) was added to each sample. Finally, samples were centrifuged at 3500 ×g for 10 min, and the MDA formed in the supernatant was determined at 532 nm with ELISA reader (Tecan, Rainbow Thermo, Austria). Tetramethoxypropane as standard was used in this test.

### 2.10. Protein carbonyl (PC) determination

Protein carbonyl content in brain mitochondrial was assessed by the spectrophotometric method. Briefly, 500 μl of 20% (w/v) trichloroacetic acid (TCA) was added to the samples (0.5 mg mitochondrial protein/ml). Afterward, the samples were located at 4 °C for 15 min. The precipitates of the samples were treated with 500 μl of 0.2% DNPH (2, 4-dinitrophenylhydrazine) and incubated for 1 hour at room temperature. 55 μl of 100 % TCA was used for precipitation of the proteins. Samples were centrifuged and washed with 1000 μl of the ethanol-ethyl acetate (1:1, v/v) mixture. Guanidine hydrochloride (200 μl, 6 M) was added to the samples and the content of the carbonyl created was calculated by reading the absorbance at 365 nm wavelength (13).

### 2.11. Determination of GSH content

The content of the GSH in the isolated mitochondria was measured using the DTNB by the spectrophotometer method. The developed yellow color by DTNB was recorded at 412 nm. GSH content in each sample was expressed as μM (16).

### 2.12. Assessment of mitochondrial toxicity

Mitochondrial toxicity was assessed as an evaluation of mitochondria function in samples by measuring the reduction of MTT (3-[4,5-dimethylthiazol-2-yl]-2,5-diphenyltetrazolium bromide with minor modification of Ghazi-Khansari *et al..* (17).

### 2.13. Statistical Analysis

Data were presented as mean ± SD. Analysis of variance (ANOVA) was performed. Newman-Keuls Multiple Comparison tests were used to determine the differences in means. All *P* values less than 0.05 were regarded as significant.

## 3. Results

The results of the asphyctic hypoxia are shown in Figure 1. The effect was dose-dependent. MgSO_4_ at doses of 500 and 600 mg/kg significantly (*P<*0.05 and *P<*0.001) prolonged the latency for death concerning the control group. At dose 600 mg/kg, MgSO_4_ showed the same activity as phenytoin which used as a positive control (*P>* 0.001). Even at the lowest tested dose, 400 mg/kg, it has prolonged survival time (30.20 ± 5.27 min) but this prolongation was not statistically significant from the control (*P>*0.05).

**Figure 1.**
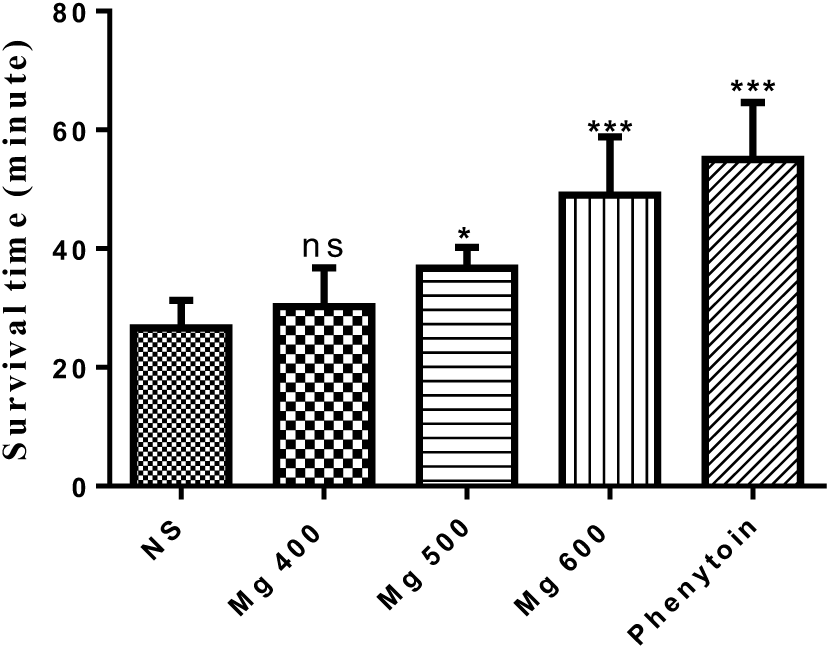
Antihypoxic activities of different doses of MgSO4 in the asphyctic hypoxia method in mice. Data are expressed as mean ± SD (n = 8)

MgSO_4_ showed good activity in the haemic model (Figure. 2). The Control group died by induced haemic hypoxia at 7.40 ± 0.46 min. MgSO_4_, at all tested doses, showed statistically significant activity concerning the control. This effect was dose-dependent. At the highest tested dose, 600 mg/kg, it has prolonged latency for death to 13.72 ± 2.88 min, which was statistically (*P<*0.01) significant from the control.

**Figure 2.**
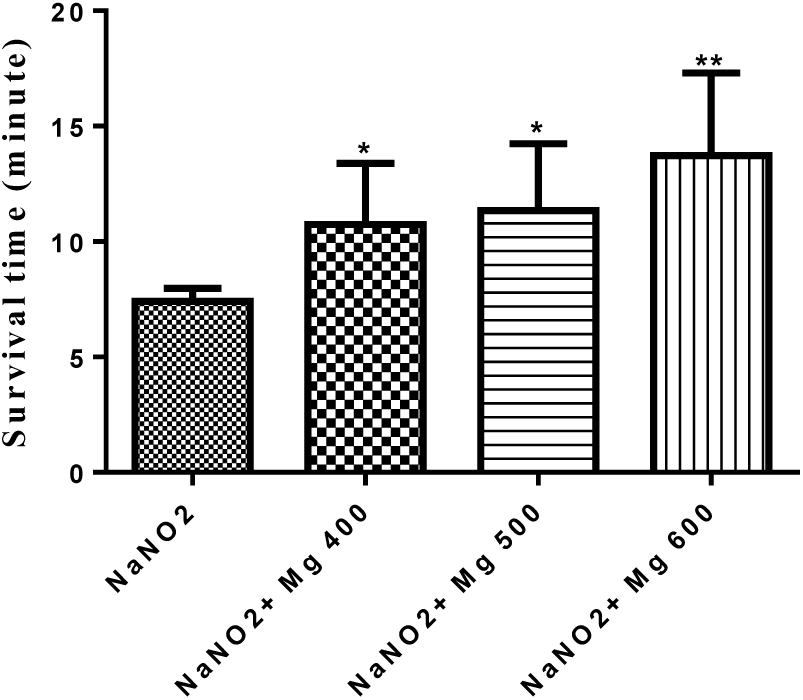
Antihypoxic activities of MgSO4 at different doses in induced-haemic hypoxia in mice. Data are expressed as mean ± SD (n = 8), *P<0.05, **P<0.01, compared to control (NaNO2).

The results of the circulatory hypoxia are shown in Figure 3. MgSO_4_ at doses of 500 and 600 mg/kg showed the most potent effect. It significantly prolonged the latency for death concerning the control group (>13.17 ± 2.06 *vs.* 6.3 ± 0.74 min, *P<*0.001). At these doses, it was showed doubles survival time and the effects were dose-dependent. At the lowest tested dose (400 mg/kg) it also kept mice alive for 7.28 ± 1.40 min but this effect was not statistically significant compared to the control (*P>*0.05).

**Figure 3.**
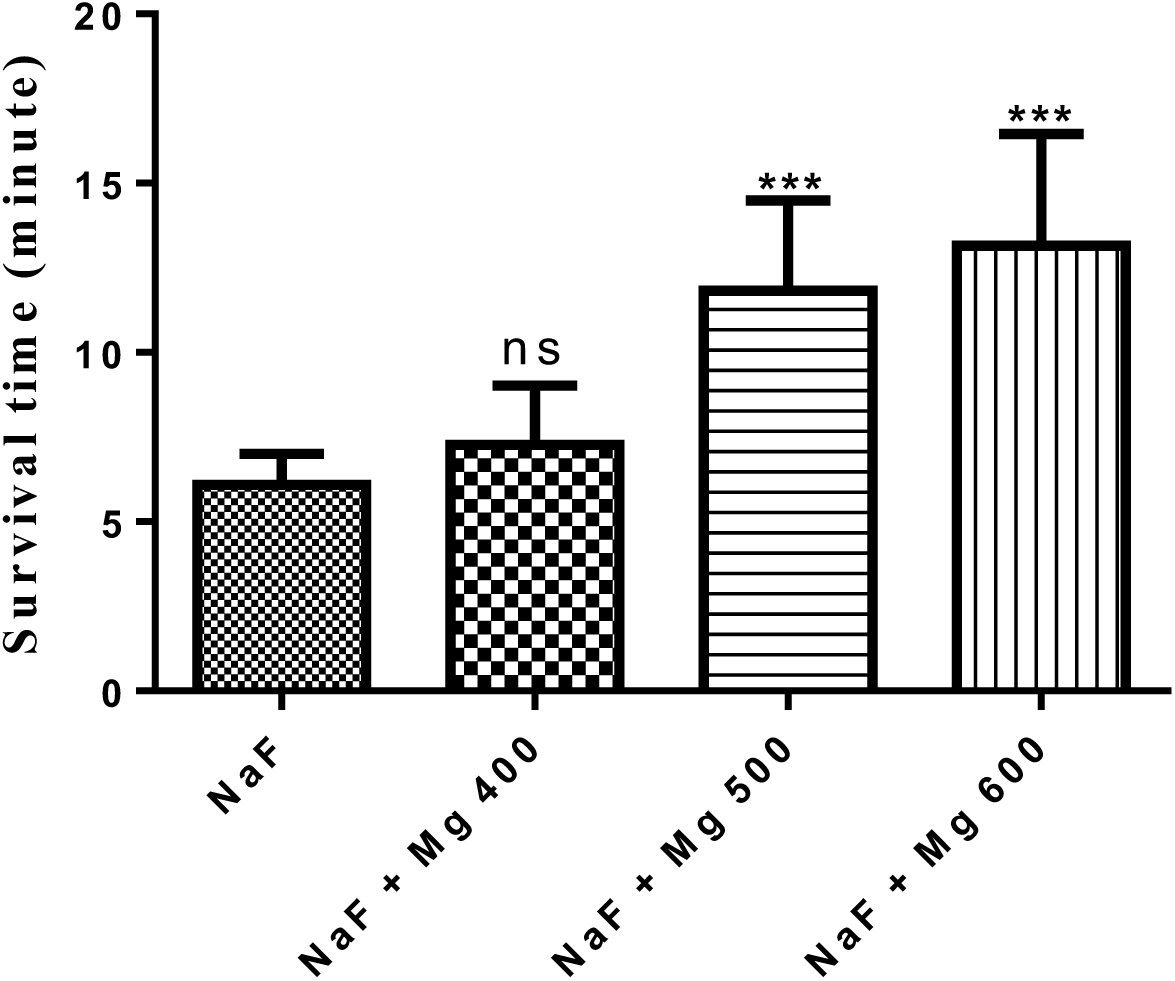
Antihypoxic activities of different doses of MgSO4 in circulatory induced-hypoxia in mice. Data are expressed as mean ± SD (n = 8), (ns, not significant; ***P<0.001, compared to control (NaF).

MDA as a marker of lipid peroxidation was assayed in the brain mitochondria of mice following different methods of induced-hypoxia. As shown in Table 1, the amount of MDA formation in mitochondria was decreased significantly (*P<*0.05) by MgSO_4_ (500 and 600 mg/kg) in haemic induced hypoxia and by MgSO_4_ (600 mg/kg) in hypoxia-induced by asphyctic and circulatory methods when compared with a control group of each.

**Table 1.** Evaluation effects of different doses of MgSO4 on lipid peroxidation, protein carbonyl formation, and glutathione content against induced hypoxia by different methods.

| Parameter → |  |  |  |  |
| --- | --- | --- | --- | --- |
| Hypoxia Type ↓ | Group | LPO<br>($\mu\text{g}/\text{mg}$ protein) | PC<br>($\mu\text{g}/\text{mg}$ protein) | GSH<br>($\mu\text{g}/\text{mg}$ protein) |
| **Asphyctic** | Con (NS) | 0.8770±0.036 | 1.320±0.097 | 0.270±0.021 |
| | Phenytoin | 0.7560±0.033 | 0.8527±0.035* | 0.1187±0.010* |
| | Mg 400 | 0.3000±0.096 | 0.7573±0.035* | 0.1813±0.045 |
| | Mg 500 | 0.6270±0.027 | 0.7883±0.028* | 0.2123±0.042* |
| | Mg 600 | 0.5020±0.210* | 0.6873±0.188** | 0.1233±0.007 |
| **Circulatory (NaF)** | Con (NS+ NaF) | 0.3468±0.038 | 1.532±0.426 | 0.1174±0.011 |
| | NaF + Mg 400 | 0.2307±0.026 | 0.4417±0.032* | 0.1077±0.005 |
| | NaF + Mg 500 | 0.2470±0.062 | 0.5247±0.074* | 0.1143±0.014 |
| | NaF + Mg 600 | 0.1470±0.010* | 0.7007±0.060 | 0.1163±0.004 |
| **Haemic ($\text{NaNO}_2$)** | Con (NS+ $\text{NaNO}_2$) | 0.6373±0.039 | 1.532 ± 0.426 | 0.148±0.022 |
| | $\text{NaNO}_2$+ Mg 400 | 0.4913±0.044 | 0.4343± 0.118* | 0.0950±0.006 |
| | $\text{NaNO}_2$+ Mg 500 | 0.2920±0.051* | 0.8483± 0.192 | 0.0846±0.004* |
| | $\text{NaNO}_2$+ Mg 600 | 0.2860±0.112* | 1.081± 0.152 | 0.0993±0.005 |

Data are presented as mean ± SD of 6 animals in each group. \* $P<0.05$ and \*\* $P<0.01$ : significantly compared with each control group.

Due to previous findings on the beneficial effects of magnesium in mitochondrial function, we decided to assess the probable benefit role of this metal on the mitochondrial antioxidant status. Mitochondrial GSH, was measured in isolated brain mitochondria after induced hypoxia with different methods. Levels of glutathione contents were significantly (*P<*0.05) increased in phenytoin and MgSO_4_ (500 mg/kg) groups compared to control in asphyctic condition and it was increased in MgSO_4_ (500 mg/kg) group in haemic (NaNO_2_) hypoxia condition compared to control (NS+ NaNO_2)_ group (Table 1). Protein carbonyl as a marker of protein oxidation in mitochondria was measured in this study. Compared to each control group (NS, NS+ NaF and NS+ NaNO_2_) PC significantly increased by treatment with Phenytoin, MgSO_4_ (400, 500 and 600 mg/kg) in asphyctic and MgSO_4_ (400 and 500 mg/kg) in circulatory and MgSO_4_ (400mg/kg) in haemic hypoxia groups (Table 1).

As shown in Figures 4 and 5, MTT reduction to Formosan, as a mitochondrial toxicity assessment, significantly (*P<*0.05) increase by MgSO_4_ (600 mg/kg) in the hypoxia-induced via asphyctic and haemic methods (44% and 66%, respectively). Mitochondria function was increased significantly (*P<*0.05) by all doses of MgSO_4_ (23%, 33%, and 67%) in circulatory hypoxia when compared to the control group (Figure 6).

**Figure 4.**
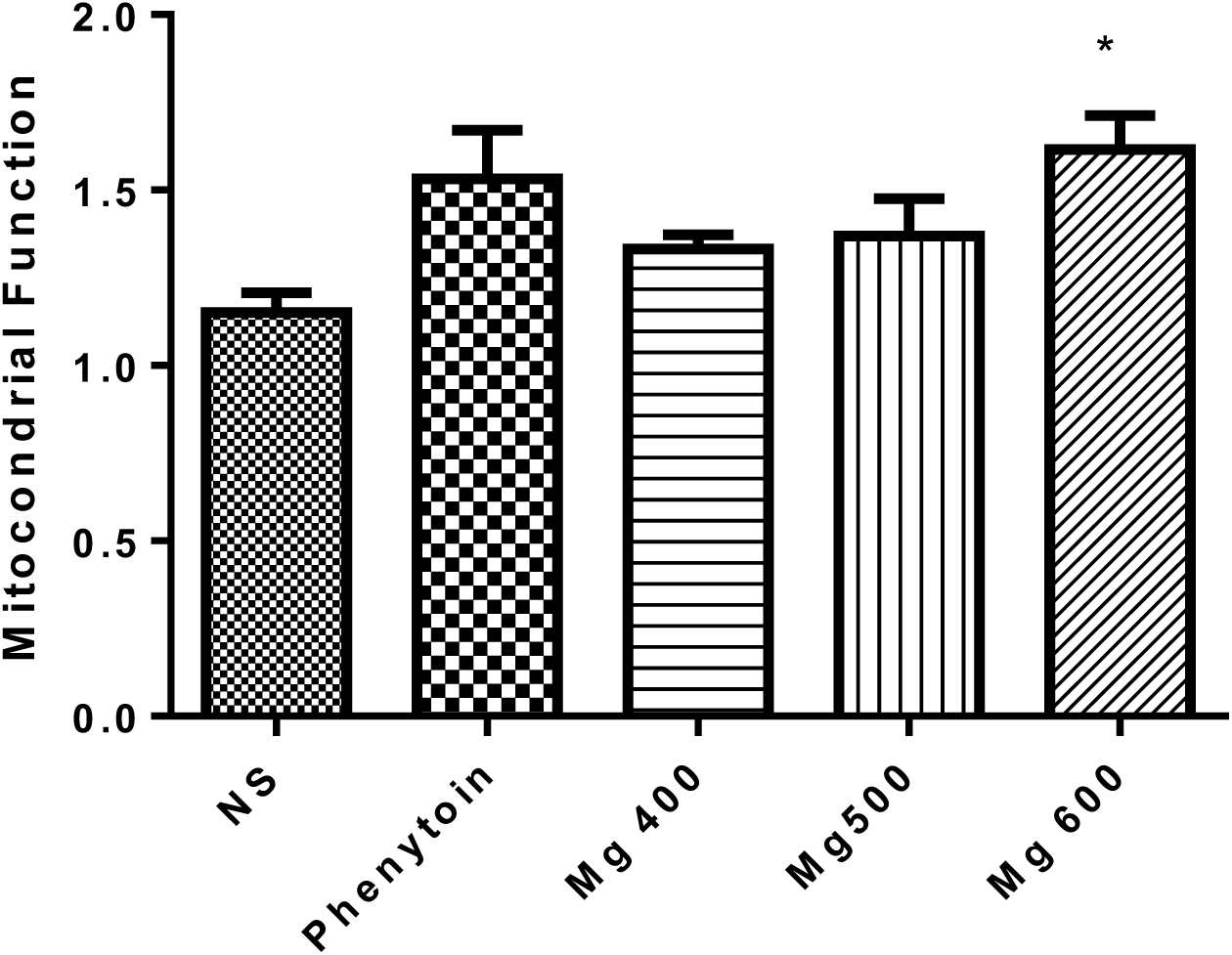
Effect of different doses of MgSO4 and phenytoin on mitochondrial function after induced hypoxia by the asphyctic method. Values represented as mean ± SD (n=8). *P < 0.05 compared with control group (NS).

**Figure 5.**
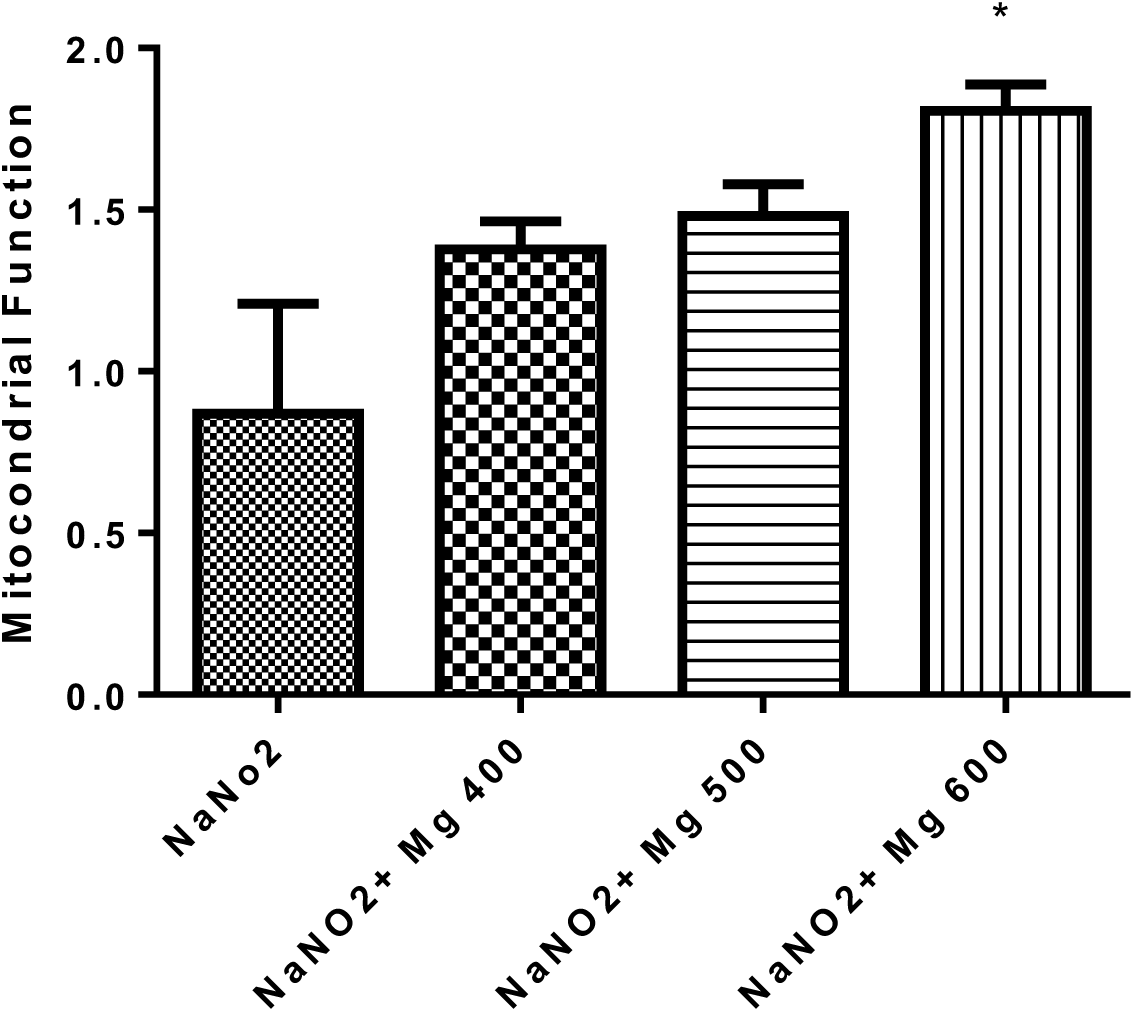
Effect of different doses of MgSO4 on mitochondrial function after induced hypoxia by Haemic (NaNO2) method. Data are presented as mean ± SD (n=8). *P<0.05 compared with control group (NaNO2).

**Figure 6.**
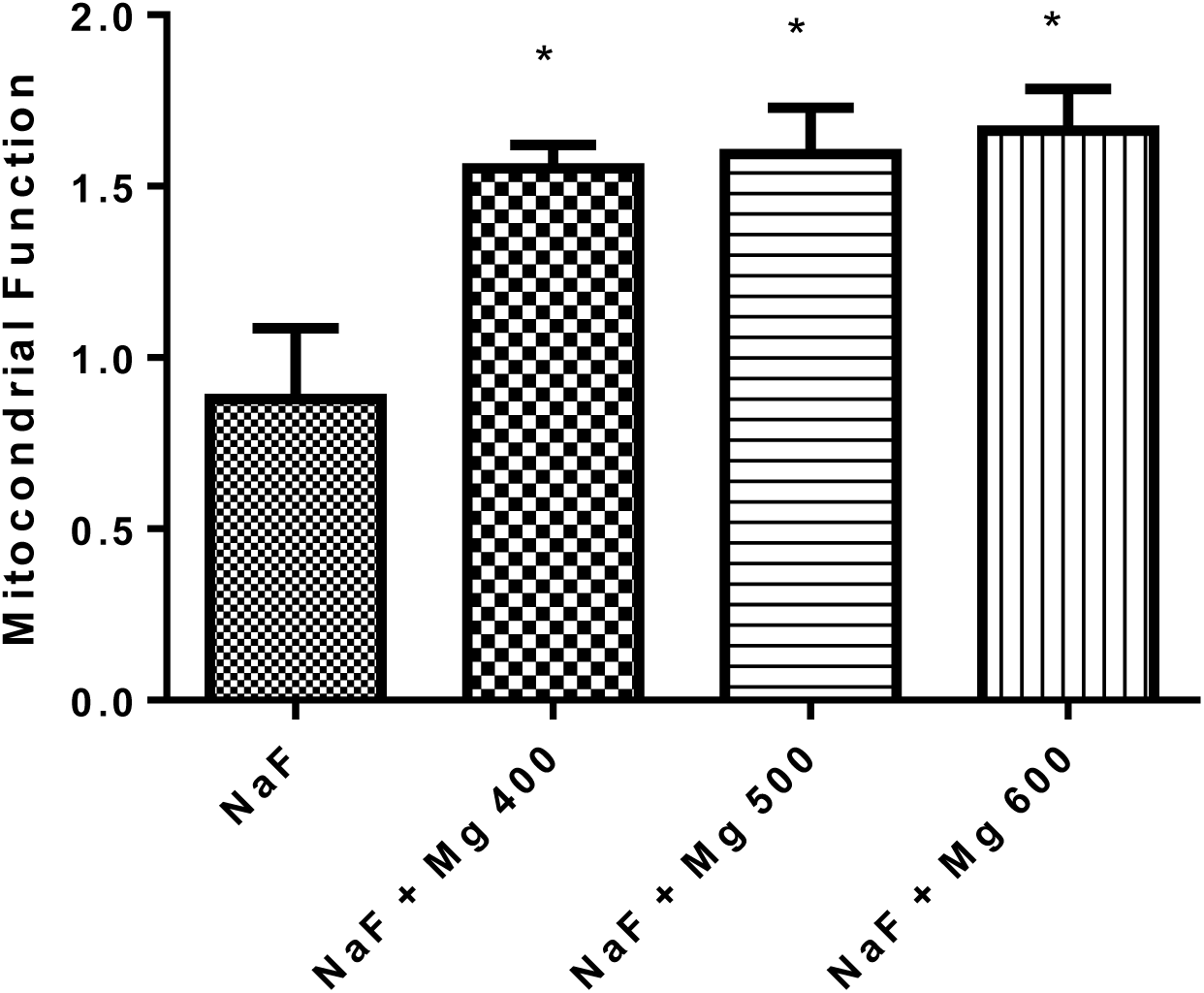
Effect of different doses of MgSO4 on mitochondrial function after induced hypoxia by Circulatory (NaF) method. Data are presented as mean ± SD (n=8). *P<0.05 compared with control group (NaF).

## 4. Discussion

This study was undertaken to evaluate how probably administration of the MgSO_4_ would reduce oxidative damage in hypoxia-induced with different methods. The results illustrated that MgSO_4_ improved oxidative stress induced by different hypoxia in a good manner.

Hypoxia produces strong physiologic stress and induces a wide range of deleterious effects at the cellular level. The brain, which consumes a large quantity of oxygen, is very vulnerable to low levels of oxygen (10) because it has a high content of polyunsaturated fatty acids, which easily undergo oxidation (18). The survival time of animals in a sealed container directly reflects the anti-hypoxic activity.

Oxygen deficiency of the brain leads to deleterious changes in the structural and functional integrity of cerebral tissue. Consequently, any drug that enables the brain to resist the consequences of ischemia or hypoxia would be of great therapeutic interest (18). During the past decades, a variety of different experimental models have been developed that could be used for testing anti-hypoxic and anti-ischemic drug effects *in vivo*.

Free radicals act as signaling species in various normal physiological processes but excessive production of these radicals causes damage to biological materials. The increased level of ROS in hypoxia is the result of the accumulation of reducing equivalents in the mitochondrial electron transport system (19). The effects of ROS can be particularly evident in certain tissues such as the brain because it consumes about one-fifth of the basal oxygen (20). Many efforts have been undertaken to develop therapies to reduce the effects of oxidative stress. Considerable evidence shows that antioxidants can exert protecting action on a variety of illnesses (21).

In this study, statistically significant anti-hypoxic activities were established in some doses of MgSO_4_ in all experimental models of induced-hypoxia in mice. The effects were dose-dependent. Nearly at all tested doses, MgSO_4_ statistically showed significant activity respect to the controls. A close relationship between oxidative metabolism and cholinergic function has been reported during the investigations of NaNO_2_ on brain metabolism (22).

Chemical hypoxia induced by the injection of NaNO_2_ will reduce the oxygen-carrying capacity of the blood by converting hemoglobin to methemoglobin. This lethal dose is injected 30 min after the MgSO_4_ treatment. Immediately after the NaNO_2_ injection, the animals are placed in small cages and the time between the injection of NaNO_2_ and cessation of respiration is recorded. Magnesium sulfate showed good activity in the haemic model (Figure 2).

There are literature data that the administration of NaF, which induces circulatory hypoxia, increases the blood histamine content, and decreases the oxygen-carrying capacity. MgSO_4_ at 600 mg/kg showed the highest activity. The mechanism of this protective action may be due in part to the antioxidant activity of MgSO_4_. Because there is no standard drug for haemic and circulatory hypoxic models, the results of this study were compared to those of control groups.

Mg is the most abundant cation with antioxidative properties that exist in more than 300 enzymatic systems. Mg is necessary for ATP activity and is vital in different processes such as current the calcium across and within the cell membranes in tissues (1, 2).

Tissue hypoxia can induce cell death by inhibition of the activity of complexes I to IV in the mitochondria respiratory chain. Moreover, the reduction of mitochondrial transmembrane potential and depletion of intracellular ATP contents can occur which finally elevates ADP/ATP ratio in the cell (23, 24). The consequence of these processes lactate dehydrogenase releases and damaging the cell membrane integrity can occur. It was demonstrated that hypoxia induces acidosis and depletion of mitochondrial energy in cells. Induce oxidative damage leads to the release of proteolytic enzymes and DNA fragmentations which processes leading to cell death (25, 26). It was reported that Mg has a beneficial role in the prevention of cell death by inhibiting calcium accumulation and improve cell metabolism (4). The present results indicated significant improvement in oxidative biomarkers when MgSO_4_ was applied to induced-hypoxic mice. The mechanism of Mg as an antioxidant defence system is not so clear and is still a matter of debate but the beneficial function of Mg in oxidative stress and damage to molecules and cells has been proven by several studies (27, 28). Hypoxia can induce ROS production that causes oxidative damage to molecules, cell organelles, and tissues. Many authors speculate that an imbalance between pro-and anti-oxidants results in an increased level of oxidative degradation of biomolecule products, such as lipid peroxidation products (29). In our study, different models of hypoxia have been used to evaluate the cellular events associated with the hypoxic injury. We induced hypoxia *via* different methods such as asphyctic, haemic (NaNO_2_), and circulatory (NaF) in mice.

In the haemic model, NaNO_2_ induces methemoglobinemia and in the circulatory model, NaF increases the blood histamine content and decreases the oxygen-carrying capacity and finally inhibits mitochondrial respiration. Demonstrated that more than 90% of Mg^2+^ in the cell is bounded to adenine nucleotides, especially ATP (4). Chemical hypoxia is distinguished by a reduction of electron carriers in mitochondrial and depletion of cell ATP leading to cell ROS formation and eventually, disruption of the plasma membrane permeability barrier with failure of cell viability (30). It has been reported that inhibition of ATP-dependent ion transporters after ATP depletion associated with hypoxic conditions results in variation in the levels of cytosolic ions. After the hypoxia onset, a rise in cytosolic free Ca^2+^ occurs which activates Ca-dependent degradative enzymes that leading to cell death. Also, chemical hypoxia can cause a large increase in free Na^+^ and H^+^ in the cell (31-33). It was stated that during hypoxia, oxygen-free radicals are produced and peroxidation of proteins and membrane lipids contribute to cell damage. Administration of MgSO_4_ to the induced-hypoxic mice showed significant attenuation of oxidative stress biomarkers compared to the control groups. Magnesium sulfate is a non-competitive and voltage-dependent antagonist of the *N*-Methyl-D-aspartate (NMDA) receptor-ion channel that may block all pathways and has been revealed to be neuroprotective in patients suffering from acute stroke and/or in various animal models of brain injury. Moreover, it was demonstrated that pretreatment with MgSO4 attenuates ROS generation and oxidative damage in the brain cells following induced hypoxia (34). Our results are compatible with other studies that have shown MgSO4 decreases oxidative damage in the brain. The observations of oxidative damage to the membrane simultaneous with mitochondrial dysfunction lend credence to the assumption that an alteration of the mitochondrial membrane is a critical step in hypoxia-induced cell death. It seems that the administration of MgSO4 prevents the fall of Mg content in the brain and attenuates the hypoxia-induced neuronal mitochondria membrane damage. These results may appear to be a conflict with previous studies indicating no effect of MgSO4 in induced hypoxic on cerebral injury in animals (35, 36).

In the present study, we demonstrated that pretreatment of mice with MgSO_4_ before hypoxia induction, not only attenuates the increased oxidative damage in the mice brain but also mitochondria function was improved by different doses of MgSO_4_ which is an important distinction of our investigation with the previous studies. It was stated that irreversible brain damage in the acute phase of hypoxia can be due to primarily necrotic changes. In regards to hypoxic tissue, diminished cytosolic Mg^2+^ is associated with increased damage and formation of ROS while increases are related to the reverse (34). The mechanism that Mg exerts its antioxidative effect is unclear but it may be associated with inhibition of iron-driven lipid peroxidation (37). Another feasibility is that Mg^2+^ blocks the excessive influx of Ca^2+^ by binding to the NMDA receptor ion channel, known as a trigger of oxygen free radical producing pathways, such as phospholipases, cyclooxygenase, and lipoxygenase (38). It has been reported that Mg^2+^ alters the neuron sensitivity to an oxidative insult (39).

Our results are agreement with prior studies that have indicated MgSO_4_ has a beneficial role in the decrease of oxidative stress biomarkers (5, 23, 34, 40). It has been demonstrated that blockade of the ionic conductance and preventing of calcium ion influx during the NMDA receptor, is the mechanism of MgSO_4_ prophylactic action in hypoxia (41). Administration of MgSO_4_ before or after the exposure to hypoxia prevents the reduction in ATP levels in cerebral tissue (42). It has been shown that acidosis can be created after hypoxia in cell tissues and consequence of that depletion of mitochondrial energy, induction of proteolytic enzymes, and oxidative damage can occur that finally result in cell death (43). Therefore, magnesium improves myocardiocytes metabolism and energy, inhibits the accumulation of calcium, and finally prevents myocardial cell death which improves delivery of oxygen to the hypoxic tissues, especially brain, and prevents the cells from hypoxic damage. Moreover, regarding different benefit results reported for Mg and magnetic Mg (^25^mg) via increasing of the creatine kinase activity (2-4 folds) and ATP level, it is obvious that Mg by enhancement of both substrate and oxidative phosphorylation pathways can increase the ATP synthesis and prevents the oxidative damage and cell death (40, 44).

It was previously confirmed that following hypoxia, ROS generation, protein, and lipid peroxidation of the neuronal cell membrane and cell membrane dysfunction were increased in the animal’s brain (34). The present changes in LPO, GSH, PC and mitochondrial function in our study supports oxidative damage potential of induced-hypoxia by different methods (Table 1 and Figures 4-6). Previous studies have shown that the generation of different free radicals such as nitric oxide, as a consequence of hypoxia, can increase in the cell (45). Therefore, complex processes such as ATP depletion, the release of excitatory amino acids, the formation of ROS, and alkoxyl radicals are the series of intracellular events due to hypoxic cell injury. Moreover, previous studies have demonstrated that Mg is involved in ATP-dependent pump activity such as Na/K-ATPase and Ca-ATPase which are sensitive to membrane lipid peroxidation (46). therefore, Mg is effective in balancing the ATP, reduction of oxidative stress, and has positive effects on cellular hypoxia. Practically, a dose of 600 mg/kg of MgSO_4_ showed the effective potential of antioxidants. Our findings showed that MgSO_4_ at dose 600 mg/kg improves hypoxia-induced oxidative stress status in mitochondria much better than other doses which may be due to more improvement of intracellular Mg levels and mitochondrial ATP.

In conclusion, this study demonstrated that during hypoxia, neuronal mitochondria oxidative damage can occur associated with a decrease of survival time in mice. Pretreatment with MgSO_4_ attenuated protein and lipid peroxidation and increased mitochondrial function in mice afflicted by different methods of hypoxia. The results of this study support the conclusion that Mg may be of worth in increasing survival time and preventing the mortality associated with asphyxiation.

## 5. Conclusion

MgSO_4_ showed a very good protective effect against hypoxia in all tested models. Specifically, it produced significant and dose-dependent benefit effects on survival time and oxidative damage in different models of induced-hypoxia.

## Acknowledgments

The authors express their appreciation to the Vice-Chancellor for Research at Mazandaran University of Medical Sciences for financial support of the current study.

